# EPIC-TRACE: predicting TCR binding to unseen epitopes using attention and contextualized embeddings

**DOI:** 10.1101/2023.06.26.546489

**Authors:** Dani Korpela, Emmi Jokinen, Alexandru Dumitrescu, Jani Huuhtanen, Satu Mustjoki, Harri Lähdesmäki

## Abstract

T cells play an essential role in adaptive immune system to fight pathogens and cancer but may also give rise to autoimmune diseases. The recognition of a peptide-MHC (pMHC) complex by a T cell receptor (TCR) is required to elicit an immune response. Many machine learning models have been developed to predict the binding, but generalizing predictions to pMHCs outside the training data remains challenging.

We have developed a new machine learning model that utilizes information about the TCR from both *α* and *β* chains, epitope sequence, and MHC. Our method uses ProtBERT embeddings for the amino acid sequences of both chains and the epitope, as well as convolution and multi-head attention architectures. We show the importance of each input feature as well as the benefit of including epitopes with only a few TCRs to the training data. We evaluate our model on existing databases and show that it compares favorably against other state-of-the-art models.

## 1 Introduction

T cells are a vital part of the adaptive immune system. To determine if an immune response is needed, T cells interact with infected, cancerous and healthy cells. Upon recognition of a target cell an immune response is elicited. This target cell recognition is based on their characterizing receptors, the T cell receptors (TCR), that bind to peptides presented by major histocompatibility complex (MHC) molecules. Thus, accurately predicting the interactions between the TCR and the peptide-MHC complex (pMHC) would be highly valuable.

The TCR consists of two chains, the *α* and the *β* chain, which both have variable regions created by somatic V(D)J-recombination. Both chains are important for the pMHC interaction and consists of three complementarity determining regions CDR1, CDR2 and CDR3. The CDR3 is the most variable region and more in contact with the peptide, whereas the CDR1 and CDR2 regions are encoded within the V gene and are more in contact with the MHC [24]. More importance has been placed on the CDR3 of the *β* chain than other parts of the TCR, which is also reflected in currently available TCR-pMHC data. However, the use of both chains and V and J gene information has been shown to improve the prediction accuracy [13, 21]. The V(D)J-recombination creates diversity both from a combinatorial effect by choosing which genes to include and a junctional effect stemming from random nucleotide insertions and deletions in the ligation process of the chosen gene segments. Together the two chains can form a vast TCR diversity with estimates ranging from 10^15^ to 10^20^, being orders of magnitudes larger than the estimated amount of cells in the human body 3.7 · 10^13^ [16]. Similarly as the TCRs, the pMHCs are very diverse. Naive estimates for pMHC diversity of one human are between 10^8^ and 10^9^ and about 10^13^ for MHC class 1 and 2, respectively [23]. In addition to the astronomical number of possible TCR-pMHC pairs, both parts show cross-reactivity, i.e. one TCR can recognize approximately 10^6^ peptides and a peptide can be recognized by many TCRs [33].

The TCR repertoire can be studied as a whole by comparing clonalities or diversities between individuals or populations [31]. The usage and evolutionary conservation of V, D and J genes have also been studied to understand the repertoires [31]. However, the underlying key concept is the TCR-pMHC binding, enabling one to understand which TCR(s) bind to which epitope(s). Many different machine learning approaches have been used to predict the TCR-pMHC binding, including clustering based methods (TCRdist [3], GLIPH [9, 10], TCRMatch [2]), decision trees (SETE [30]), random forests (TCRex [8], epiTCR [22]), and Gaussian processes (TCRGP [13]). Recently, as more data has become available, many different data-intensive deep learning approaches have been proposed (ERGO [29, 28], ImRex [21], TITAN [32], NetTCR [15, 20], DeepTCR [27], TCRAI [35], TCRconv [14], TEINet [12]). Still, the complexity of the problem and the quality, amount and imbalance of the available data cause challenges for developing methods that generalize to TCRs and pMHC not included in the training data. Many of the methods use only parts of the TCRs, e.g. only CDR3s or only the *β* chain [30, 21, 32, 20, 27, 22, 12]. Moreover, most available prediction tools use epitopes only as categorical features, omitting the amino acid sequence altogether [3, 9, 10, 2, 30, 8, 13, 27, 35, 14]. This effectively leads to inability to predict binding for epitopes outside the training data. Furthermore, already limited training data has to be filtered out when training epitope-specific predictors, as there are not sufficient amounts of data per epitope. On the contrary, the ability to predict for unseen peptides would facilitate the prediction of cognate peptides to disease-associated orphan TCRs.

In this work we present a new deep learning model, EPIC-TRACE, that utilizes ProtBERT [4] based contextualized encodings of the amino acid sequences of the peptide and the TCR as well as multi-head attention and convolutions to achieve accurate and robust predictions. We primarily focus on predicting TCR-pMHC interactions for peptides that are not included in the training data (so-called unseen epitope task). As input to the EPIC-TRACE model we use the CDR3, V and J genes of both chains (whenever available) and the peptide sequence together with its corresponding MHC allele. We show that utilizing information about all available parts of the TCR-pMHC complex as input features in our model leads to best predictive performance. Furthermore, we show that including peptides that may have only a few interacting TCRs in the training data improves the performance on the unseen epitope task and demonstrate how the model can be used as an in silico peptide screening method. Finally we show that our model performs better or comparable to recent models across a variety of prediction tasks.

## 2 Methods

### 2.1 Data

TCR-pMHC discovery relies mostly on the use of pMHC-multimers, which are restricted to relatively few pMHCs compared to a vast amount of possible T cells screened for recognition. Thus, the current TCR-pMHC data is skewed to have far more unique TCRs than pMHCs. This skewed data makes the TCR-pMHC prediction task harder. We collected our data of positive TCR-pMHC pairs from two databases: VDJdb [1] and IEDB [18]. Since both VDJdb and IEDB have much less MHC class II datapoints, we filtered the data to contain only MHC class I datapoints and further required the host to be from human. For a fair comparison we define our base dataset 𝒟_*αβ,β*_, which we subsample or extend as explained later in the corresponding experiments. For each datapoint in 𝒟_*αβ,β*_, we required the following information: the amino acid sequence of the *β* chain CDR3 region, the epitope amino acid sequence, and information about the *β*V, *β*J and MHC genes at any precision, i.e. the full-length amino acid sequence of the TCR might not be available. Dataset 𝒟_*αβ,β*_ contains only datapoints with sufficient *β* information, to which we also add information about *α* chain, when available. In Results Section 3 we use 𝒟_*αβ,β*_ as described above unless specified otherwise.

We unified the notation for all V and J genes and discarded datapoints with non-functional genes according to the IMGT [17, 25, 26, 5, 6]. We ensured that all CDR3s are in canonical form by adding missing anchor position residues (C and F/W) and if not possible we discarded the data-point. Datapoints that only differed in precision of gene information were filtered out by keeping only the most precise. The numbers of unique feature values of all datasets are shown in Suppl. Table S1. We note that the three most frequent epitopes, i.e. epitopes with most associated TRCs, make up more than a fourth of all datapoints in our IEDB + VDJdb based datasets.

### 2.2 Prediction tasks

Because of the paired nature of the data and the challenges due to the data imbalance, the TCR-pMHC prediction is more suitable to be expressed as four separate tasks. An important distinction is whether to test for epitopes contained in the training data (seen epitopes) or the converse, test for unseen epitopes. Methods treating the epitope as categorical label cannot naturally predict for unseen epitopes. This distinction is very important as it precisely defines the difficulty of the problem. Furthermore, following [29] the tasks can similarly be divided in terms of seen or unseen TCRs, resulting in the following three tasks: TCR-Peptide Pairing 1 (TPP1) where both TCR and epitope parts of a test datapoint are seen in training data but in different pairs, TPP2 where the epitope is seen in training but TCR is unseen, and TPP3 where neither the TCR nor the epitope is seen in training. To complete the task definitions we add TPP4, where the TCR is seen in training data but the epitope is unseen.

The different tasks correspond to different biological questions. TPP2 seeks to answer if a TCR repertoire has T cells targeting given epitope(s), e.g. SARS-COV-2 or HIV, such that we have data on those epitopes in training. In the case our training data contains neither the epitopes nor the TCRs, the task changes to TPP3, which is arguably the most general and interesting task. Even though all tasks are relevant, currently only the two first tasks (TPP1 and TPP2) can be solved with reasonable performance. However, as individuals have generally very little overlap in their TCR repertoires the unseen TCR tasks (TPP2 and TPP3) are more generally applicable. In addition, TCRs in the current databases, such as IEDB and VDJdb, are mostly specific to a single epitope, i.e., the TCRs appear as a pair to only one epitope. This means that the amount of positive datapoints for the TPP4 task is very low and makes it unfeasible to test with. Thus, we focus our experiments primarily to the TPP3 task, but we also include TPP2 experiments as a comparison. For the TPP4 evaluation we use an external dataset that we describe later.

### 2.3 Cross-validation and performance metrics

Following the common practice in the field, the performance of our model was evaluated using 10-fold cross-validation that was repeated five times. In each cross-validation fold we split the data to train and test sets and extract part of the train set for validation used for early stopping. We report the performance measures as the mean and standard error of the mean across the five cross-validation runs. We use the area under the receiver operating characteristics (AUROC) and the average precision (AP) metrics.

As the data attained from the databases contained only positive pairs, the negative data had to be generated. Optimally, one would use experimentally derived negatives, such as those from the 10x study [7], but these are not generally available for all epitopes. The generation of negative data by shuffling the positive datapoints is established in the field and gives a more reliable estimate of model performance compared to usage of external TCR datasets [21]. The negatives are generated separately for every train (+ validation) and test set in each cross-validation fold to ensure a larger amount of negatives to shuffle epitopes in train. Importantly, this also restricts data leakage from test to train for the corresponding task. We generated the negative data by shuffling TCRs (CDR3, V and J for both chains) with epitopes (epitope and MHC) such that the new datapoint was not in the set of positive datapoints. The randomly generated datapoint was determined negative if any part (CDR3, V or J gene) of either chain was different to the positive TCRs of the epitope. As epiTRC, TITAN and ImRex only consider the *β* chain and the epitope, the above definition can create some CDR3*β*-Epitope pairs that have both positive and negative labels. To ensure a fair comparison with epiTCR, TITAN and ImRex, we created a second version of cross-validation splits of 𝒟_*αβ,β*_, where we determined the negatives such that at least the CDR3*β* had to differ. In both settings we did not allow duplicate datapoints. We created five times as many negatives as positives for each epitope as in [20, 19]. This means that not only has the test set the ratio of 1:5 but also any individual epitope. For most frequent epitopes, there are not enough TCRs to create enough negatives by shuffling. In these cases we discarded positive datapoints randomly to maintain the correct ratio. In order to comply with the given TPP task definitions, the splits were created separately for each task.

As the majority of the datapoints belong to a small amount of most frequent epitopes, we balanced the amount of epitopes and datapoints in each fold for TPP3. More specifically, we ordered the epitopes in descending frequency order and then randomly assigned a fold index for *k* = 10 consecutive epitopes at a time. Importantly, this also assures that all folds have both frequent and less frequent epitopes. The TCRs are more evenly distributed and thus the cross-validation folds for TPP2 can be done simply by choosing TCRs to the splits. If any of the epitopes in the test set is not present in train, extra negatives were added to the train to obtain the TPP2 constraint (and naturally the 1:5 ratio cannot be retained for those epitopes).

### 2.4 EPIC-TRACE model

Our model utilizes the full TCR-pMHC information available and is designed to predict interaction between a TCR and an pMHC, i.e., a binary classification problem.

#### TCR features

A TCR is defined as TCR = (*β*_V_, *β*_J_, *β*_CDR3_, *α*_V_, *α*_J_, *α*_CDR3_), where 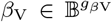 and 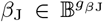 are one-hot encoded vectors indicating the V and J genes in the *β* chain, 𝔹 = {0, 1}, and *g*_*β*V_ and *g*_*β*J_ denote the numbers of V and J genes (similarly for the *α* chain, 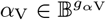 and 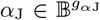). Variable 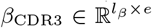 consists of two parts: *(i)* contextualized information about the CDR3 region of the *β* chain that is obtained from the pre-trained ProtBERT language model (size *l*_*β*_ × 1024) [4], and *(ii)* one-hot encoded CDR3 region. These are concatenated to form feature representation of size *l*_*β*_ × *e*, where *l*_*β*_ denotes the length of the CDR3 region that is further padded to CDR3 maximum length *l* × *e*. If the V and J gene information is available for the *β* chain, the full-length TCR *β* amino acid sequence is constructed and embedded with ProtBERT. For full-length TCRs only the CDR3 region positions are extracted from the ProtBERT embedding and stored in *β*_CDR3_. This is done as we use the V and J genes as separate inputs, and as shown by [14] the contextualized CDR3 captures the essential features for classification. If the full TCR cannot be constructed, only the CDR3 region is embedded with ProtBERT and no further extraction is done. If the *α* chain is available, 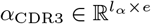 is defined similarly.

#### Epitope-MHC features

The epitope-MHC complex is defined as pMHC = (Epitope, MHC), where Epitope 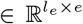 is obtained by concatenating the ProtBERT embedding and the one-hot encoding of the epitope sequence (and subsequent padding to maximum length), *l*_*e*_ is the length of epitopes, 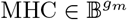 is the one-hot encoded vector of the MHC allele, and *g*_*m*_ is the number of alleles.

#### Output labels

We formulate our model using three separate binary output labels *y* = (*y*_*α*_, *y*_*β*_, *y*_*αβ*_), where 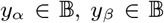, and *y*_*αβ*_ ∈ 𝔹. If only the *β* chain is available, then *y*_*α*_ and *y*_*αβ*_ are considered as missing (similarly if only *y*_*α*_ is available). If both *α* and *β* chains are available, then *y*_*αβ*_ defines the binding and *y*_*α*_ and *y*_*β*_ are considered missing. The prediction problem is then defined with datapoints (TCR_*n*_, pMHC_*n*_, *y*_*n*_), where *n* ∈ {1, …, *N* }, and *N* is the number of positive and negative datapoints.

#### Architecture

Our architecture utilizes convolutions, multi-head self-attentions, learnable linear embeddings, non-linearities and dropouts, and the model contains three output heads corresponding to the cases when only *β*, only *α*, or both *β* and *α* chains are available. Overview of the model is shown in Fig. 1. The representations of the CDR3 regions (*β*_CDR3_ and *α*_CDR3_) and the epitope (Epitope) are first processed with 1-D convolutions. Epitope convolution is concatenated separately with the CDR3 convolution of the *β* and *α* chains (if available). Multi-head attention is used to identify the important interacting features separately for (*β*_CDR3_, Epitope) and (*α*_CDR3_, Epitope) pairs. Learnable linear embedding is trained for the one-hot encoded V and J genes from both chains (*β*_V_, *β*_J_, *α*_V_, *α*_J_) as well as for the MHC allele (MHC), which are then concatenated with the outputs of the attentions. The *β* and *α* chains are processed with the multilayer perceptrons (linear and ReLu) separately as well as together (whenever both chains are available) and passed through sigmoidal activation to make the predictions *ŷ*_*β*_ ∈ [0, 1], *ŷ*_*α*_ ∈ [0, 1] and *ŷ*_*αβ*_ ∈ [0, 1] corresponding to the three different cases. Details of the neural network architecture are shown in Suppl. Section S2.

**Figure 1.**
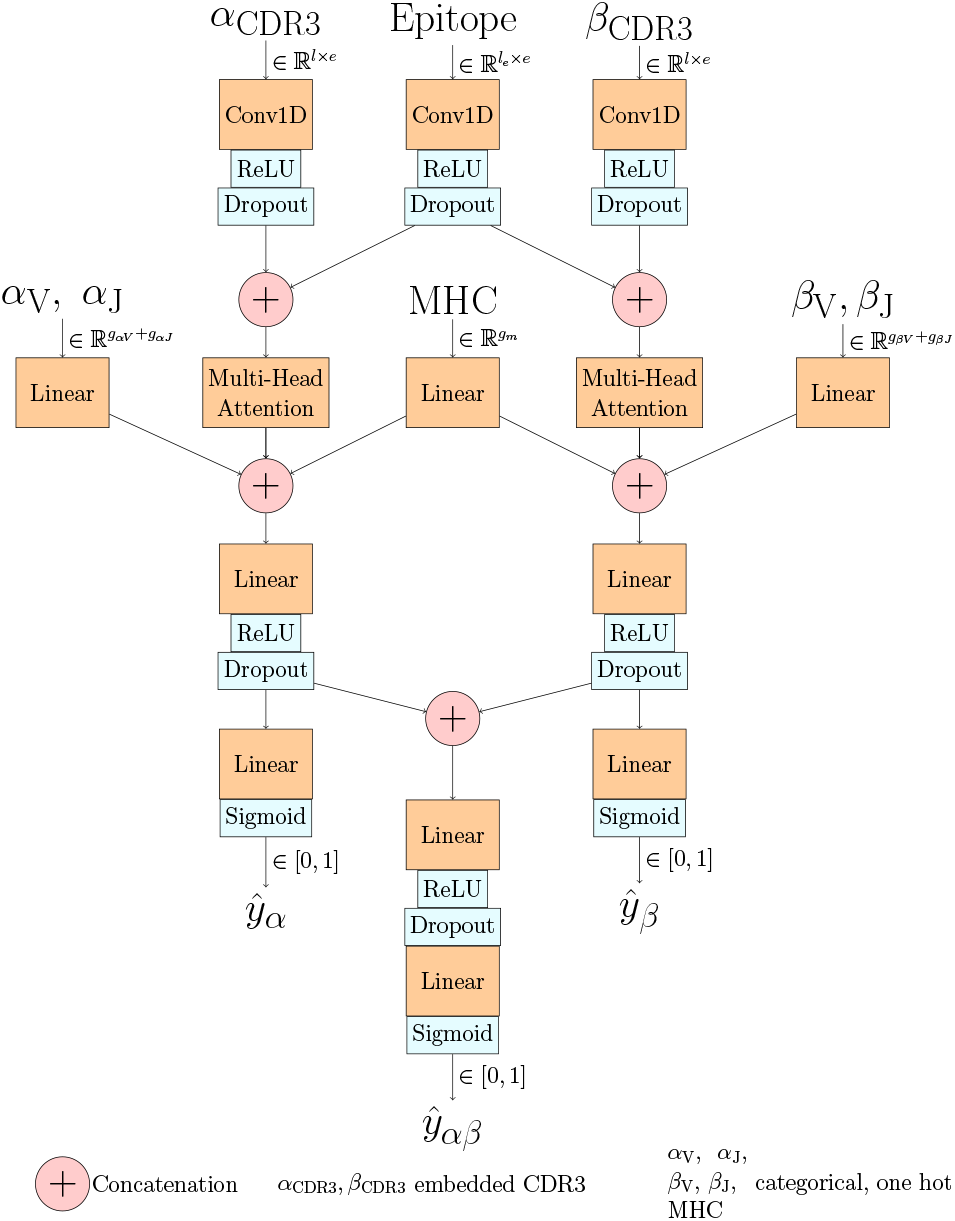
Architecture of the EPIC-TRACE model. The model takes as input either or both of the TCR chains and their corresponding V and J genes in addition to the epitope/peptide sequence and the MHC allele. The amino acid sequences are contextualized with ProtBERT and gene inforation is provided as one hot vectors. Depending on the availability of the chains one of the models three output heads is used for the prediction.

#### Model training

We trained our model by maximizing the logarithm of the Bernoulli likelihood or equivalently the negative binary cross-entropy

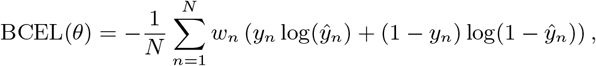

where *w*_*n*_ is the weight for the *n*^th^ datapoint. We weighted the positive datapoints five times higher than the negatives. We controlled for over-fitting by using early stopping based on the average precision on the validation set, and the model parameters giving the highest validation score were used. Following the main training we used stochastic weight averaging (SWA) [11] for 20 epochs. We used different learning rates for training models for the TPP2 and TPP3 tasks in both the main training and the SWA sampling, 0.0001 and 0.001, respectively. In addition we used exponential learning rate scheduler for the main training for TPP3.

## 3 Results

### 3.1 Choice of validation set, and per epitope scores

We first set out to investigate the effect of the validation set (used for early stopping) on the test performance. We compared two different ways to generate the validation set: *(i)* naive random sample of datapoints from the train set, and *(ii)* creating unseen epitope validation by choosing datapoints by epitopes from the train set. The comparison was made only for the TPP3 task, where epitope is unseen, as the unseen epitope validation is not sensible for seen epitope tasks TPP1 or TPP2.

The random validation set had slightly better performance compared to the unseen epitope validation (see Table 1), perhaps because that leaves more distinct epitopes in the training set. The unseen epitope relation is present between both train-validation (TPP3 or TPP4) and train-test (TPP3). However, the epitope distributions in validation and test are naturally distinct for the TPP3 task. Due to this inherent epitope covariate shift (as a result of very few epitopes in the current data) a representative validation set is hard to construct. Because the random validation is better representing the other tasks and also resulted in slightly better performance for the TPP3, we chose to use the random validation in all following experiments.

**Table 1:** Comparison of validation strategies. The model was trained with either randomly selected validation data or validation data that had a unseen epitope (TPP3 or TPP4) relation to the remaining train set. Validation data was used for early stopping and selecting model parameters before SWA. Reported values are the mean and standard error of five 10-fold cross-validation runs.

|  | TPP3 AUROC | TPP3 AP |
| --- | --- | --- |
| Random validation | $0.691 \pm 0.008$ | $0.291 \pm 0.005$ |
| Unseen epitope validation | $0.688 \pm 0.010$ | $0.285 \pm 0.008$ |

Due to the highly imbalanced data the joint prediction accuracy measures (AUROC and AP) are dominated by the epitopes with most data-points. Therefore, we quantified the per epitope scores for the TPP2 and TPP3 tasks. We observe that the epitopes with more datapoints have a higher score on average on the TPP2 task (Fig. 2), which is logical as there are more datapoints for those epitopes to train on. To better characterize the trend explained by the number of datapoints for an epitope in the TPP2 task, we binned the per epitope scores and calculated the bin averages (Fig. 3a). The AUROC scores seem to slightly increase as the number of datapoints increases. On the other hand, we observe that the number of datapoints per epitope does not affect the performance on the TPP3 task as expected (Suppl. Fig. S1), since by the TPP3 definition the datapoints for a specific epitope are not included in the training and, thus, only affect the number of test datapoints. Overall, prediction accuracies vary across epitopes, which can be due to the currently available data for that epitope or underlying biophysical reasons.

**Figure 2.**
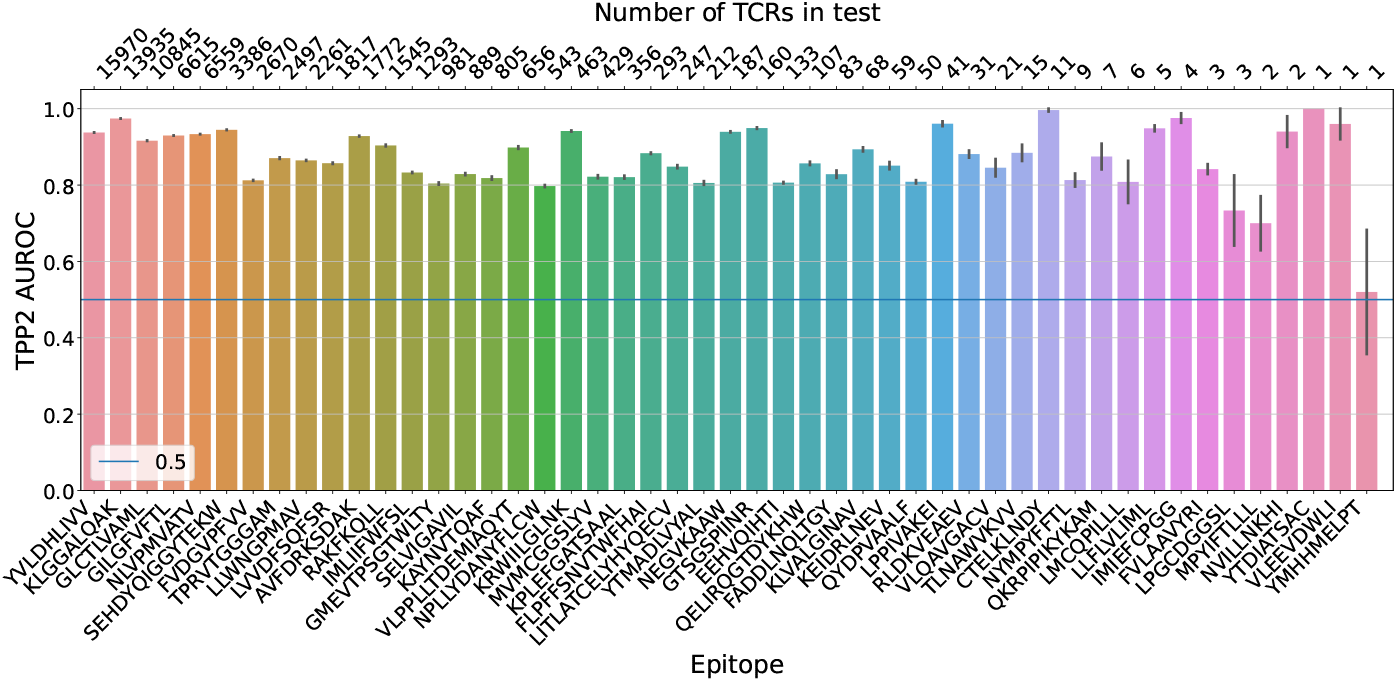
Per epitope AUROC values for the TPP2 task. Epitopes were sampled logarithmically to include epitopes with varying number of TCRs. Top *x*-axis shows the number of positive datapoints for each epitope (bottom *x*-axis). The vertical axis shows the mean of five 10-fold cross-validations runs together with the standard error.

**Figure 3.**
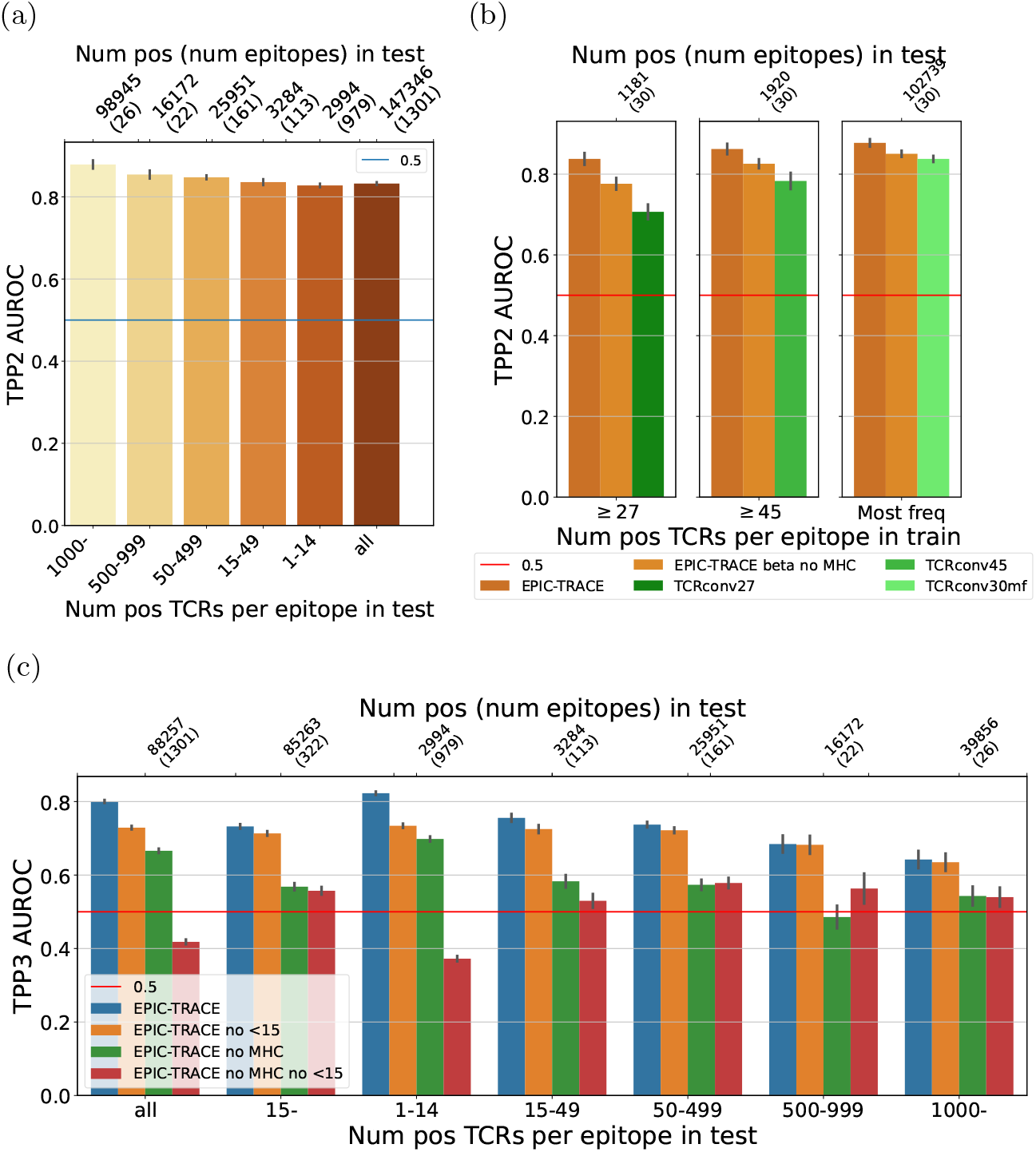
a) Average of per epitope AUROC values for epitopes binned by datapoint frequency in the TPP2 task. Epitopes were binned to five bins according to the number of positive datapoints to assess frequency based trend. b) Comparison to TCRconv. TCRconv was trained on three subsets of 30 epitopes from the 𝒟_*αβ,β*_ dataset and compared to EPIC-TRACE trained on full 𝒟_*αβ,β*_ folds either with all or reduced input features. The *y*-axis shows average per epitope AUROC values of frequency binned epitopes with standard error. c) Comparison of models trained with all datapoints or by discarding epitopes with less than 15 TCRs from training for TPP3. Models were trained with or without MHC information. The *y*-axis shows the average per epitope AUROC with standard error.

Furthermore, we investigated the effect of a distance between epitopes in the training and test sets. This was done by quantifying the minimum (Levenshtein) edit distance between an epitope in the test set and the epitopes in the training set. In addition to the simple edit distance we tried a BLOSUM based and a BERT embedding based distance. Similarly as in [21] we observe that the per epitope scores seems to slightly decrease when the minimum edit distance to the training set increases (Suppl. Fig. S2).

### 3.2 Input feature contribution

To study which input features are important we conducted an ablation study and trained the model using different features. We studied the performance gain of using the full length TCRs (long context) when possible for creating ProtBERT embeddings from which the CDR3 part is extracted, compared to using always only the CDR3 region as input for the ProtBERT model. In addition, we trained the models with or without the categorical V, J and MHC information. Models were trained separately for TPP2 and TPP3 tasks. The results are presented in Table 2 and discussed below.

**Table 2:** Effect of input features. The model was trained on 𝒟_*αβ,β*_ using different subsets of the input features. Here CDR3 and long in parenthesis denote the context used for the ProtBERT embeddings and VJ and MHC denote if the respective categorical features were used. We also compared the model on 𝒟_*αβ,α,β*_ that contains also datapoints that have only the *α* chain but not the *β* chain (row 8). Reported values are the mean of the five 10-fold cross-validation runs together with the standard error.

|  | TPP2 AUROC | TPP2 AP | TPP3 AUROC | TPP3 AP |
| --- | --- | --- | --- | --- |
| 1. $\alpha\beta$ (CDR3) | $0.830 \pm 0.000$ | $0.574 \pm 0.000$ | $0.513 \pm 0.008$ | $0.179 \pm 0.003$ |
| 2. $\alpha\beta$ (CDR3) + VJ | $0.891 \pm 0.000$ | $0.665 \pm 0.001$ | $0.548 \pm 0.007$ | $0.192 \pm 0.004$ |
| 3. $\alpha\beta$ (CDR3) + MHC | $0.837 \pm 0.000$ | $0.583 \pm 0.000$ | $0.611 \pm 0.002$ | $0.243 \pm 0.002$ |
| 4. $\alpha\beta$ (CDR3) + VJ + MHC | $0.897 \pm 0.000$ | $0.676 \pm 0.000$ | $0.692 \pm 0.007$ | $0.289 \pm 0.006$ |
| 5. $\alpha\beta$ (long) | $0.888 \pm 0.000$ | $0.663 \pm 0.000$ | $0.528 \pm 0.008$ | $0.191 \pm 0.004$ |
| 6. $\alpha\beta$ (long) + MHC | $0.893 \pm 0.000$ | $0.674 \pm 0.000$ | $0.682 \pm 0.010$ | $0.284 \pm 0.007$ |
| 7. $\alpha\beta$ (long) + VJ + MHC | **$0.906 \pm 0.000$** | **$0.698 \pm 0.000$** | $0.691 \pm 0.008$ | $0.291 \pm 0.005$ |
| 8. $\alpha\beta$ (long) + VJ + MHC [$\mathcal{D}_{\alpha\beta,\alpha,\beta}$] | **$0.906 \pm 0.000$** | $0.691 \pm 0.001$ | **$0.693 \pm 0.008$** | **$0.294 \pm 0.007$** |

Interestingly, the two tasks benefited in different magnitudes of the different features. The VJ gene information given either as categorical features or as part of the context to the ProtBERT embeddings was more important for the TPP2 task, while the MHC information was more important for the TPP3 task (Table 2). Even though the vast majority of the datapoints had either “HLA class 1” or “HLA*02:01” as their MHC information the MHC feature showed to be important. For the TPP3 task, the performance without the MHC information is much lower than when using it, even if VJ information is used. The results are logical when comparing the MHC importance between the tasks. In the TPP3 case the MHC information can be included in the training data, and thus some information about the pMHC complex can be directly used in test predictions. On the other hand, in the TPP2 task, most of the datapoints for one epitope share the same MHC information and thus this information becomes redundant, which explains the lower improvement. Expectedly, both tasks had best performance when using both VJ and MHC information. When using both VJ and MHC information the gene-gene preferences can be explicitly modelled, which could explain synergistic improvement on the TPP3 task.

To investigate the importance of the TCR chains, we evaluated the EPIC-TRACE model on a reduced dataset (𝒟_*αβ*_), where every datapoint has necessary information of both chains available. With 𝒟_*αβ*_ we required that TCRs in the test sets differed on both *α* and *β* CDR3s from TCRs in the train. Similarly, we required both CDR3s to differ when determining negative pairs. We trained our model utilizing only either chain and with both chains. Results are shown in Table 3. As expected, using both chains outperforms the models that were trained on only the *α* or *β* chain. However, when using only either chain the performances are very similar and, somewhat surprisingly, the *α* chain provides even a slightly better performance. We note that the performances on the reduced dataset 𝒟_*αβ*_ are worse than on the full dataset 𝒟_*αβ,β*_ due to a smaller sample size.

**Table 3:** Comparison of TCR chains. The model was trained with either or both of the TCR chains on a more stringent dataset 𝒟_*αβ*_, where each datapoint contains both chains. Reported values are the mean of the five 10-fold cross-validation runs together with the standard error.

| Used chain(s) | TPP2 AUROC | TPP2 AP | TPP3 AUROC | TPP3 AP |
| --- | --- | --- | --- | --- |
| $\alpha\beta$ | $0.767 \pm 0.001$ | $0.505 \pm 0.001$ | $0.541 \pm 0.008$ | $0.204 \pm 0.006$ |
| $\beta$ | $0.721 \pm 0.001$ | $0.441 \pm 0.001$ | $0.539 \pm 0.004$ | $0.202 \pm 0.005$ |
| $\alpha$ | $0.725 \pm 0.001$ | $0.442 \pm 0.001$ | $0.537 \pm 0.005$ | $0.206 \pm 0.004$ |

Lastly, we combined our base dataset (i.e., 𝒟_*αβ,β*_ that contains both *αβ* and *β* datapoints) with datapoints containing only the *α* chain (i.e., 𝒟_*αβ,α,β*_). This could not be done with the other models that are compared in this paper as they require *β* chain (ERGO-II) or can only utilize either of the chains (epiTCR, TITAN and ImRex). The combination was done by adding the new datapoints to the training sets leaving the test sets the same and comparable. Adding the *α* datapoints increased the TPP3 performance but lowered the AP on the TPP2, see row 8 Table 2

### 3.3 Increasing the amount of unique epitopes improves generalisation

To investigate how the number of unique epitopes in the training data affects the two tasks (TPP2 and TPP3), we evaluated the model with two settings: *(i)* we included all epitopes in the cross-validation (i.e., the same standard cross-validation as above), and *(ii)* we discarded the epitopes with less than 15 TCRs from training. These settings were also extended to test sets such that the test set either included or excluded the less frequent epitopes. These low frequency epitopes comprise approximately 75 % of the (1301) epitopes but only 2994 of the 147346 datapoints. In earlier work low frequency epitopes have been discarded from the data: e.g. epitopes with less than 15 TCRs were excluded in TITAN [32], and epitopes with less than 10 were excluded in TEInet [12]. The performance scores for the two tasks and the two different settings are shown in Table 4. When testing on all epitopes, we observe an increase in the performance for the TPP3 task when the low frequency epitopes are included in the training data. Interestingly, when testing on more frequent epitopes (i.e., epitopes that have at least 15 TCRs), we observe increased performance in the TPP3 task but not in the TPP2 task. To further investigate the effect of low frequency epitopes on the low frequency and the more frequent epitopes separately, we calculated the average per epitope scores for the different settings. Fig. 3c shows that including the low frequency epitopes in training significantly improves the results on the TPP3 task. This is especially apparent for the low frequency epitopes in the test set. Overall, the result in Fig. 3c show that utilizing the low frequency epitopes in training is beneficial for generalisation. We note that the low frequency epitopes are associated to many HLA alleles that are not present in the data of the more frequent epitopes. Since the MHC information improves the results on the TPP3 task as shown in Section 3.2, we wanted to confirm that it is indeed the addition of different epitope sequences that improves the result, not just the addition of MHC alleles. To confirm that, we trained our model without the MHC information in the same two settings. Fig. 3c shows that including low frequency epitopes in the training data results in a similar performance improvement even when the EPIC-TRACE model is trained without the MHC information, thus supporting our hypothesis.

**Table 4:** Comparison of models trained with all data or by discarding low frequency epitopes (i.e., epitopes with less than 15 TCRs) from training. Reported values are the mean of five 10-fold cross-validation runs together with the standard error. NA indicates that the setting is not consistent with the TPP2 task definition.

| Test data | Train data | TPP2 AUROC | TPP2 AP | TPP3 AUROC | TPP3 AP |
| --- | --- | --- | --- | --- | --- |
| All | All | $0.906 \pm 0.000$ | $0.698 \pm 0.000$ | $0.691 \pm 0.008$ | $0.291 \pm 0.005$ |
| | <15 Discarded | NA | NA | $0.681 \pm 0.009$ | $0.280 \pm 0.008$ |
| $\geq 15$ | All | $0.907 \pm 0.000$ | $0.701 \pm 0.000$ | $0.686 \pm 0.008$ | $0.285 \pm 0.005$ |
| | <15 Discarded | $0.907 \pm 0.000$ | $0.701 \pm 0.001$ | $0.680 \pm 0.009$ | $0.278 \pm 0.006$ |

### 3.4 Comparisons to other methods

Next we compared our method to other state of the art models that treat the epitope as an amino acid sequence. We compared against ERGO-II [29, 28], TITAN [32], ImRex [21], and epiTCR [22]. We used again the dataset 𝒟_*αβ,β*_ and exactly the same cross-validations data splits for all methods. The results in Table 5 show that our model outperforms epiTCR, TITAN and ImRex by a large margin, and performs consistently better than ERGO-II on both tasks. One reason to the difference can be that epiTCR, TITAN and ImRex only utilize the *β* chain and the *β*-CDR3, respectively, compared to our model and ERGO-II utilising all available information. Performance of all models remained consistent when using the different definitions for negative datapoints (see Suppl. Table S2).

**Table 5:** Comparison to previous methods. EPIC-TRACE, ERGO-II and epiTCR are evaluated on five 10-fold cross-validation runs, whereas TITAN and ImRex are evaluated on only one of the 5 cross-validations due to long training time. Reported values are the mean of the five 10-fold cross-validation runs together with the standard error.

|  | TPP2 AUROC | TPP2 AP | TPP3 AUROC | TPP3 AP |
| --- | --- | --- | --- | --- |
| EPIC-TRACE (our) $[\mathcal{D}_{\alpha\beta,\alpha,\beta}]$ | **0.906** ± 0.000 | 0.691 ± 0.001 | **0.693** ± 0.008 | **0.294** ± 0.007 |
| EPIC-TRACE (our) | **0.906** ± 0.000 | **0.698** ± 0.000 | 0.691 ± 0.008 | 0.291 ± 0.005 |
| ERGO-II | 0.895 ± 0.002 | 0.659 ± 0.007 | 0.675 ± 0.007 | 0.274 ± 0.004 |
| epiTCR | 0.793 ± 0.000 | 0.581 ± 0.000 | 0.515 ± 0.001 | 0.183 ± 0.001 |
| TITAN 700 | 0.786 | 0.454 | 0.577 | 0.204 |
| ImRex 700 | 0.697 | 0.420 | 0.519 | 0.178 |

We also compared our model against a state of the art model that uses epitopes as class labels, TCRconv [14]. Since the number of unique epitopes in the dataset originally used for TCRconv is in the order of tens, we trained TCRconv separately with three subsets of 30 epitopes from the 𝒟_*αβ,β*_, stratified according to the number of TCRs per epitope in the train set (i.e., epitopes with ≥27, ≥45, or ≥780 TCRs in the train set, the last one presenting the most frequent epitopes). This was done for a more fair comparison as opposed to using hundreds of epitopes. For a more detailed description of the comparison see Supplementary material Section S1. Fig. 3b shows that EPIC-TRACE performs better on all three subsets with both the full model and the reduced model (only *β* chain and no MHC). As expected, the more frequent epitopes receive a better mean AUROC score than the less frequent epitopes for both EPIC-TRACE and TCRconv. Importantly, the difference between TCRconv and EPIC-TRACE increases when the epitope frequency decreases, showcasing the advantage of using the epitope amino acid sequence. We also tested EPIC-TRACE against TCRconv on the most abundant epitopes using both *α* and *β* sequences. This is a setting where methods that treat epitopes as class labels are strongest. We observed that TCRconv can achieve a comparable performance in this setting (see Suppl. Table S3), but as discussed above, TCRconv or other similar tools cannot make prediction for any other epitopes than those in the training data.

### 3.5 Prediction of yeast display data

Next we demonstrate how EPIC-TRACE can be used to screen epitopes for disease-associated TCRs — a computational task that is notoriously difficult but would have tremendous potential e.g. in understanding disease pathogenesis. Recently, [34] identified five orphan TCRs that are associated with ankylosing spondylitis (AS) as well as acute anterior uveitis (AAU) and used yeast display library screening followed by subsequent validation to identify 26 HLA-B*27:05 restricted shared self-peptides and microbial peptides that activated the five AS- and AAU-derived TCRs. Here we demonstrate that machine learning methods are starting to reach sufficient accuracy to complement, and eventually replace, the laborious yeast display library screening. We used the five experimentally validated TCRs and 26 epitopes, altogether 81 HLA-B*27:05 restricted TCR-peptide pairs, as positive datapoints, and created negative datapoints by assigning 2000 randomly selected HLA-B*27:05 restricted epitopes from IEDB to the five TCRs. EPIC-TRACE model trained with VDJDB + IEDB dataset 𝒟_*αβ,α,β*_ performed poorly on the yeast display dataset (AU-ROC 0.485). This was also the case for ERGO-II (AUROC 0.195). This prediction task is challenging because all the datapoints have the same HLA and V alleles, meaning that the distinction has to be made purely by peptide and CDR3 sequences. Therefore, we added 10% of the 81+2000 datapoints (corresponding on average to 8 positive datapoints) as part of the training data to provide EPIC-TRACE model information about the AS and AAU associated TCR-peptide pairs, and then evaluated the model performance on the remaining 90% of the data (unseen epitope, TPP4). To account for the different epitopes we split the 2081 datapoints from the yeast display dataset in 10 parts by epitopes and evaluated the performance for each part in the train. The average AUROC and AP scores of the 10 parts were 0.784 and 0.281 respectively. The recall and number of true positives against the number of best scoring test datapoints in the first split are shown in Fig. 4. This analysis shows that by utilizing as little as 10%, or on average 8 positive datapoints, the performance of the model in the yeast display library task is at least moderately good.

**Figure 4.**
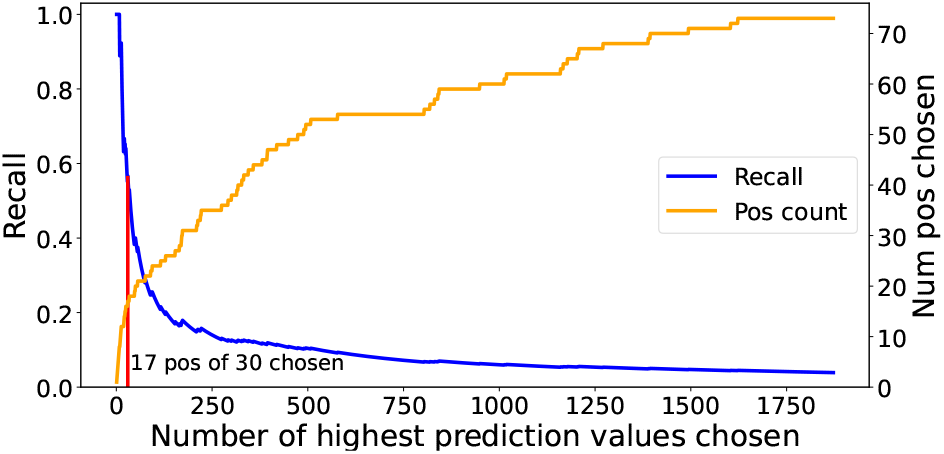
Recall and positive count by number of chosen datapoints. EPIC-TRACE was trained with the IEDB + VDJDB data and 10% (of which 8 positive datapoints) of the generated yeast display data. The test data contained 73/1872 positive datapoints.

## 4 Discussion

Here, we have presented EPIC-TRACE, a novel method for predicting TCR-pMHC binding using the full TCR information together with the peptide amino acid sequence and MHC allele. We showed that the seen and unseen epitope tasks behave differently and have different importance for the used input features. It is apparent that current data mostly obtained with the use of pMHC-multimers is very imbalanced and leads to difficulties to generalize to the full TCR-pMHC space. More specifically, the unseen epitope task remains very hard for state of the art methods. We showed that specificity to some epitopes is easier to predict than to others, which results in varying predictive performance across epitopes. Although the simple minimum edit distance to train set in the TPP3 case explained the general difficulty, it is not accurate enough to be used as an estimate for prediction accuracy for any specific epitope. An estimate of the reliability of the prediction would be very useful for both the seen and unseen tasks.

Surprisingly, we found out that epitopes with fewer datapoints were easier to predict for in the TPP3 task. The reason could be a difference in the underlying methods to obtain the data. Furthermore, the development and use of new TCR-pMHC sequencing methods increase the throughput and quality of the data. Especially important is that the amount of distinct epitopes increases, even if these epitopes are not associated to many TCRs, thus also the unseen epitope task becomes more feasible to solve.

## Acknowledgements

We acknowledge the computational resources provided by the Aalto Science-IT project.

## References

[1] Dmitry V Bagaev et al. “VDJdb in 2019: database extension, new analysis infrastructure and a T-cell receptor motif compendium”. In: Nucleic Acids Research 48.D1 (2020), pp. D1057–D1062.

[2] William D Chronister et al. “TCRMatch: Predicting T-cell receptor specificity based on sequence similarity to previously characterized receptors”. In: Frontiers in immunology 12 (2021), p. 640725.

[3] Pradyot Dash et al. “Quantifiable predictive features define epitope-specific T cell receptor repertoires”. In: Nature 547.7661 (2017), p. 89–93. doi: https://doi.org/10.1038/nature22383.

[4] Ahmed Elnaggar et al. “ProtTrans: Towards Cracking the Language of Lifes Code Through Self-Supervised Deep Learning and High Performance Computing”. In: IEEE Transactions on Pattern Analysis and Machine Intelligence (2021), p. 1–1. doi: 10.1109/TPAMI.2021.3095381.

[5] Géraldine Folch and Marie-Paule Lefranc. “The human T cell receptor beta diversity (TRBD) and beta joining (TRBJ) genes”. In: Experimental and clinical immunogenetics 17.2 (2000), p. 107– 114.

[6] Géraldine Folch and Marie-Paule Lefranc. “The human T cell receptor beta variable (TRBV) genes”. In: Experimental and clinical immunogenetics 17.1 (2000), p. 42–54.

[7] 10 Genomics. “A New Way of Exploring Immunity–Linking Highly Multiplexed Antigen Recognition to Immune Repertoire and Phenotype”. In: Tech. rep (2019).

[8] Sofie Gielis et al. “Detection of enriched T cell epitope specificity in full T cell receptor sequence repertoires”. In: Frontiers in immunology (2019), p. 2820.

[9] Jacob Glanville et al. “Identifying specificity groups in the T cell receptor repertoire”. In: Nature 547.7661 (2017), p. 94– 98. doi: https://doi.org/10.1038/nature22976.

[10] Huang Huang et al. “Analyzing the Mycobacterium tuberculosis immune response by T-cell receptor clustering with GLIPH2 and genome-wide antigen screening”. In: Nature biotechnology 38.10 (2020), p. 1194–1202.

[11] Pavel Izmailov et al. “Averaging weights leads to wider optima and better generalization”. In: arXiv preprint arXiv:1803.05407 (2018).

[12] Yuepeng Jiang, Miaozhe Huo, and Shuai Cheng Li. “TEINet: a deep learning framework for prediction of TCR–epitope binding specificity”. In: Briefings in Bioinformatics 24.2 (2023), bbad086.

[13] Emmi Jokinen et al. “Predicting recognition between T cell receptors and epitopes with TCRGP”. In: PLoS computational biology 17.3 (2021), e1008814.

[14] Emmi Jokinen et al. “TCRconv: predicting recognition between T cell receptors and epitopes using contextualized motifs”. In: Bioinformatics 39.1 (2023), btac788.

[15] Vanessa Isabell Jurtz et al. “NetTCR: sequence-based prediction of TCR binding to peptide-MHC complexes using convolutional neural networks”. In: BioRxiv (2018), p. 433706.

[16] Daniel J Laydon, Charles RM Bangham, and Becca Asquith. “Estimating T-cell repertoire diversity: limitations of classical estimators and a new approach”. In: Philosophical Transactions of the Royal Society B: Biological Sciences 370.1675 (2015), p. 20140291. doi: http://doi.org/10.1098/rstb.2014.0291.

[17] Marie-Paule Lefranc et al. “IMGT unique numbering for immunoglobulin and T cell receptor variable domains and Ig superfamily V-like domains”. In: Developmental & Comparative Immunology 27.1 (2003), p. 55–77.

[18] Swapnil Mahajan et al. “Epitope specific antibodies and T cell receptors in the immune epitope database”. In: Frontiers in immunology (2018), p. 2688.

[19] Pieter Meysman et al. “Benchmarking solutions to the T-cell receptor epitope prediction problem: IMMREP22 workshop report”. In: ImmunoInformatics 9 (2023), p. 100024.

[20] Alessandro Montemurro et al. “NetTCR-2.0 enables accurate prediction of TCR-peptide binding by using paired TCRα and β sequence data”. In: Communications biology 4.1 (2021), p. 1– 13.

[21] Pieter Moris et al. “Current challenges for unseen-epitope TCR interaction prediction and a new perspective derived from image classification”. In: Briefings in Bioinformatics 22.4 (2021), bbaa318. doi: https://doi.org/10.1093/bib/bbaa318.

[22] My-Diem Nguyen Pham et al. “epiTCR: a highly sensitive predictor for TCR–peptide binding”. In: Bioinformatics 39.5 (Apr. 2023). btad284. ISSN: 1367-4811. doi: 10.1093/bioinformatics/btad284. eprint: https://academic.oup.com/bioinformatics/article-pdf/39/5/btad284/50204900/btad284.pdf.URL: https://doi.org/10.1093/bioinformatics/btad284.

[23] Kenneth L Rock, Eric Reits, and Jacques Neefjes. “Present yourself! By MHC class I and MHC class II molecules”. In: Trends in immunology 37.11 (2016), p. 724–737.

[24] Markus G Rudolph and Ian A Wilson. “The specificity of TCR/pMHC interaction”. In: Current opinion in immunology 14.1 (2002), p. 52–65.

[25] Dominique Scaviner and Marie-Paule Lefranc. “The human T cell receptor alpha joining (TRAJ) genes”. In: Experimental and Clinical Immunogenetics 17.2 (2000), p. 97–106.

[26] Dominique Scaviner and Marie-Paule Lefranc. “The human T cell receptor alpha variable (TRAV) genes”. In: Experimental and clinical immunogenetics 17.2 (2000), p. 83–96.

[27] John-William Sidhom et al. “DeepTCR is a deep learning framework for revealing sequence concepts within T-cell repertoires”. In: Nature communications 12.1 (2021), p. 1–12.

[28] Ido Springer, Nili Tickotsky, and Yoram Louzoun. “Contribution of t cell receptor alpha and beta cdr3, mhc typing, v and j genes to peptide binding prediction”. In: Frontiers in immunology 12 (2021).

[29] Ido Springer et al. “Prediction of specific TCR-peptide binding from large dictionaries of TCR-peptide pairs”. In: Frontiers in immunology (2020), p. 1803.

[30] Yao Tong et al. “SETE: Sequence-based Ensemble learning approach for TCR Epitope binding prediction”. In: Computational Biology and Chemistry 87 (2020), p. 107281.

[31] Sebastiaan Valkiers et al. “Recent advances in T-cell receptor repertoire analysis: bridging the gap with multimodal singlecell RNA sequencing”. In: ImmunoInformatics (2022), p. 100009.

[32] Anna Weber, Jannis Born, and Marıa Rodriguez Martınez. “TITAN: T-cell receptor specificity prediction with bimodal attention networks”. In: Bioinformatics 37.Supplement 1 (2021), p. i237–i244.

[33] Linda Wooldridge et al. “A single autoimmune T cell receptor recognizes more than a million different peptides”. In: Journal of Biological Chemistry 287.2 (2012), p. 1168–1177.

[34] Xinbo Yang et al. “Autoimmunity-associated T cell receptors recognize HLA-B* 27-bound peptides”. In: Nature (2022), p. 1–7.

[35] Wen Zhang et al. “A framework for highly multiplexed dextramer mapping and prediction of T cell receptor sequences to antigen specificity”. In: Science Advances 7.20 (2021), eabf5835.

