## Supplementary material for "EPIC-TRACE: predicting TCR binding to unseen epitopes using attention and contextualized embeddings"

#### S1 TCRconv comparison

In the first setting we trained and evaluated TCRconv with the 30 least frequent epitopes that had at least 27 TCRs (TCRconv27) or 45 TCRs (TCRconv45) in the train set and with the 30 most frequent epitopes (TCRconv30mf) (See Epi27, Epi45 and Epi780, respectively Table S1). In this setting TCRconv was trained utilizing only the $\beta$ chain as all datapoints did not contain $\alpha$ chain information. TCRconv used the full contextualised chain (similarly as EPIC-TRACE) when available. EPIC-TRACE was trained on the full $\mathcal{D}_{\alpha\beta,\beta}$ cross-validation train sets but tested only on the 30 corresponding epitopes such that the testing data was identical. This was done both using all available features and comparing using a reduced model with only the $\beta$ chain and without the MHC information.

In the second setting the cross-validation was stratified such that the train set contained always at least 45 (positive) TCRs with both chains available for each epitope. Here both methods were trained utilizing both chains and using the full context if available (precise V and J genes). To show the benefit of using single chain datapoints in addition to the $\alpha\beta$ datapoints we trained EPIC-TRACE also by adding the $\alpha$ and $\beta$ datapoints corresponding to the epitopes already in the train set. Lastly we added all other datapoints to the train set of EPIC-TRACE. From Table S3 we can see that when training on the same data TCRconv performs better on both AUROC and AP (rows 1 and 4) . However, as corresponding single chain datapoints are added to the train set, both scores are improved and EPIC-TRACE performs better in terms of AUROC (row 2). Further adding other epitopes does not increase the performance on the epitopes in test (row 3), which could be due to the already sufficient amount of data for the specific epitopes in the test set.

### S2 Model specifics

First 1D convolutional layers ($\alpha$, $\beta$ and Epitope) have 100 output channels a kernel size of 7 and stride 1. The convolutional layers are followed by Dropout with a rate of 0.2. The Multi-head self attention modules ($\alpha$-Epitope and $\beta$-Epitope) have 5 heads and use a dropout rate of 0.2. All allele information ($\beta_V, \beta_J, \alpha_V, \alpha_J$ and MHC) is embedded with the linear layers to dimension 8. The following linear layers after concatenation ($\alpha$ and $\beta$ branches) have dimensions $2924 \times 54$ and a dropout rate of 0.45. These are followed by the output heads where the $\alpha\beta$ output head first has linear layer of dimension $108 \times 36$ with dropout rate 0.45 before the final linear layer and the sigmoid activation.

### S3 Figures and Tables

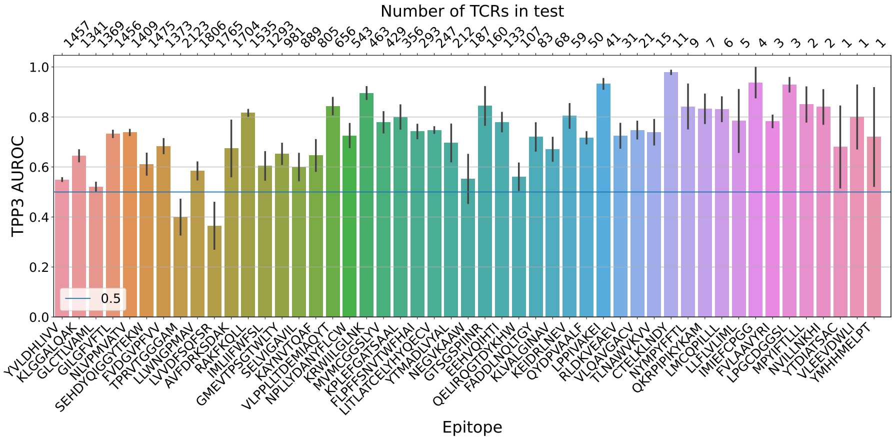


Figure S1: Per epitope AUROC values for the TPP3 task. Epitopes were sampled logarithmically to include epitopes with varying number of TCRs. Top $x$-axis shows the number of positive datapoints for each epitope (bottom $x$-axis). The vertical axis shows the mean of five 10-fold cross-validations runs together with the standard error.

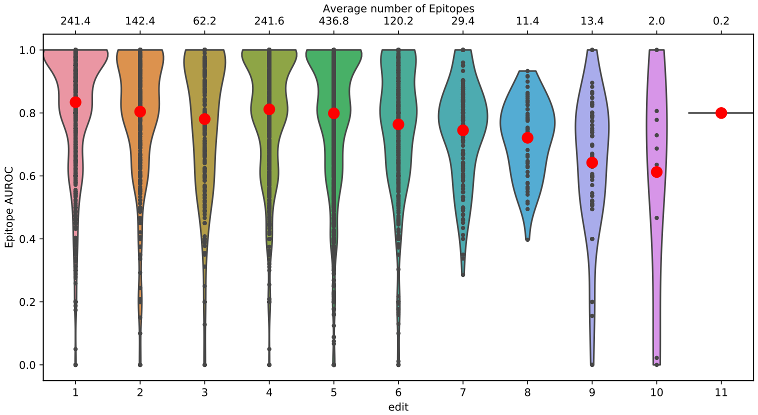


Figure S2: Violin plot of AUROC scores grouped by minimum edit distance to train dataset. The red dots are (unweighted) averages of the scores for the given minimum edit distance.

Figure S3:

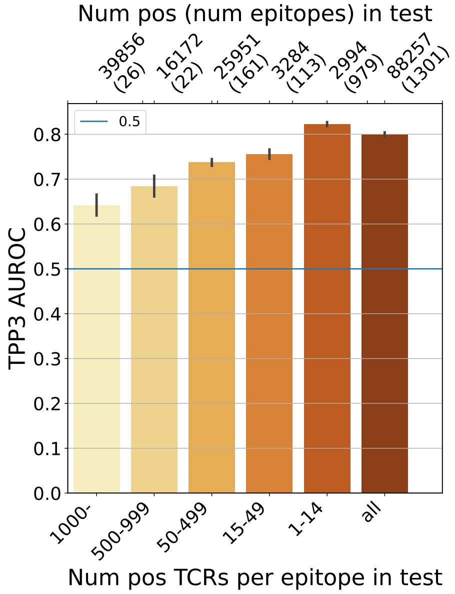


Figure S4: Average of per epitope AUROC values for epitopes binned by datapoint frequency in the TPP3 task. Epitopes were binned to five bins according to the number of positive datapoints to assess frequency based trend.

Table S1: Dataset description. Number of unique features for the five IEDB + VDJdb based datasets and the external yeast display dataset ($\mathcal{D}_{\text{YD}}$).

| | datapoints | $\beta_{\text{CDR3}}$ | $\beta_V$ | $\beta_J$ | $\beta_{\text{long}}$ | $\alpha_{\text{CDR3}}$ | $\alpha_V$ | $\alpha_J$ | $\alpha_{\text{Long}}$ | MHC | Epitope |
|---|---|---|---|---|---|---|---|---|---|---|---|
| $\mathcal{D}_{\alpha\beta,\beta}$ | 147346 | 118896 | 110 | 17 | 77022 | 19744 | 98 | 65 | 19765 | 69 | 1301 |
| $\mathcal{D}_{\alpha\beta,\alpha,\beta}$ | 169792 | 118907 | 110 | 17 | 77022 | 34692 | 106 | 65 | 28390 | 69 | 1307 |
| $\mathcal{D}_{\alpha\beta}$ | 28377 | 21266 | 106 | 17 | 21110 | 19744 | 98 | 65 | 19765 | 54 | 919 |
| $\mathcal{D}_{\alpha\beta\geq 50}$ | 25178 | 18817 | 104 | 17 | 18727 | 17532 | 91 | 65 | 17606 | 15 | 32 |
| Epi27 | 1181 | 1088 | 86 | 15 | 533 | 247 | 65 | 52 | 201 | 9 | 30 |
| Epi45 | 1920 | 1786 | 85 | 15 | 1110 | 270 | 59 | 55 | 260 | 13 | 30 |
| Epi780 | 102739 | 80538 | 102 | 17 | 58043 | 15629 | 84 | 64 | 15710 | 9 | 30 |
| $\mathcal{D}_{\text{YD}}$ | 81 | 4 | 1 | 2 | 4 | 5 | 1 | 5 | 5 | 1 | 26 |

Table S2: Negative similarity and method comparison. All models were evaluated on two datasets were *(i)* negatives were generated from positives such that any change in the TCR (V, J or CDR3 or either chain) from a positive pair was defined as a plausible negative (TCR-similarity), and *(ii)* at least a difference in the $\beta$CDR3 region was required to generate a negative from the positives ($\beta_{\text{CDR3}}$-Similarity). TITAN an ImRex are only evaluated on the first of the five cross-validations. The difference between the negative generation methods is small and $< 1\%$ of the negatives have a corresponding positive $\beta$CDR3-Epitope pair.

| | | TPP2 AUROC | TPP2 AP | TPP3 AUROC | TPP3 AP |
|---|---|---|---|---|---|
| TCR-Similarity | $\alpha\beta$ (CDR3) + VJ +MHC | 0.897 ± 0.000 | 0.676 ± 0.000 | 0.692 ± 0.007 | 0.289 ± 0.006 |
| | ERGO-II | 0.895 ± 0.002 | 0.659 ± 0.007 | 0.675 ± 0.007 | 0.274 ± 0.004 |
| | TITAN | 0.454 | 0.786 | 0.204 | 0.577 |
| | ImRex | 0.420 | 0.697 | 0.178 | 0.519 |
| | epiTCR | 0.793 ± 0.000 | 0.581 ± 0.000 | 0.515 ± 0.001 | 0.183 ± 0.001 |
| $\beta_{\text{CDR3}}$-Similarity | $\alpha\beta$ (CDR3) + VJ +MHC | 0.896 ± 0.000 | 0.678 ± 0.001 | 0.700 ± 0.008 | 0.294 ± 0.007 |
| | ERGO-II | 0.894 ± 0.002 | 0.653 ± 0.007 | 0.687 ± 0.007 | 0.278 ± 0.005 |
| | TITAN | 0.453 | 0.786 | 0.200 | 0.566 |
| | ImRex | 0.423 | 0.699 | 0.178 | 0.522 |
| | epiTCR | 0.792 ± 0.000 | 0.580 ± 0.000 | 0.516 ± 0.001 | 0.184 ± 0.000 |

Table S3: $\alpha\beta$ data ablation study. EPIC-TRACE was trained and tested on epitopes with at least 50 $\alpha\beta$ TCRs (32 epitopes). This was compared to a setting where single chain datapoints for the corresponding epitopes were added to the train set (row 2) and a setting where full data including other epitopes were included in the train set (row 3).

| | AUROC | AP |
|---|---|---|
| $\alpha\beta$ | 0.843 ± 0.003 | 0.641 ± 0.006 |
| $\alpha\beta \cup$ corresponding $\alpha$ and $\beta$ | 0.853 ± 0.002 | 0.668 ± 0.004 |
| Full data | 0.853 ± 0.002 | 0.668 ± 0.005 |
| TCRconv | 0.848 ± 0.002 | 0.676 ± 0.006 |
